# Regulation of trophic architecture across spatial scales in a major river network

**DOI:** 10.1101/317644

**Authors:** Eric Harvey, Florian Altermatt

## Abstract

Moving beyond species count data is an essential step to better understand the effects of environmental perturbations on biodiversity and ecosystem functions, and to eventually better predict the strength and direction of those effects. Here, coupling an integrative path analysis approach with data from an extensive countrywide monitoring program, we tested the main spatial, environmental and anthropogenic drivers of change in stream macroinvertebrate trophic structure along the entire Swiss Rhine river catchment. Trophic structure was largely driven by inherent altitudinal variation influencing and cascading to regional scaled factors such as land use change and position in the riverine network, which, in turn, transformed local habitat structure variables. Those cascading effects across scales propagated through the biotic community, first affecting preys and, in turn, predators. Our results illustrate how seemingly less important factors can act as essential transmission belts, propagating through direct and indirect pathways across scales to generate the specific context in which each trophic group will strive or not, leading to characteristic landscape wide variations in trophic community structure.

## INTRODUCTION

River ecosystems constitute iconic examples of spatial complexity with complex regional scale vertical structures (from upstream to downstream; the river network) constraining organism and energy movement^1–5^, but also strong localized horizontal interactions with the terrestrial matrix influencing local habitat characteristics through changes in cross-ecosystem subsidy^6–8^. The shape of river networks, which all follows the same geometric scaling properties^2^, has been shown to influence biological community dynamics and local species richness patterns^3,5,9–12^. However, recent studies have found that the relative importance of the regional river network and local habitat characteristics is somewhat context-dependant as a function of species traits (e.g., dispersal mode) and location-specific conditions such as terrestrial land-use and biotic interactions^13–15^. Although those studies tend to emphasize the importance of considering both local and regional factors to understand variations in aquatic community, total explanatory power remains generally low^16^. In that context, the use of well-defined functional or trophic groups each including taxonomically different but functionally similar taxa could improve explanatory power by generating groups of taxa with more uniform response to specific environmental or spatial characteristics^17^.

In addition, current approaches tend to focus on the relative importance of regional versus local factors to identify the dominant drivers while totally ignoring the inherent structure of interdependences among regional and local factors leading to a general loss of explanatory power^16, 18–23^. The often-assumed dichotomy between regional and local factors generally erodes when considering the mechanisms behind those effects^22,24^. For instance, many regional factors, such as altitude, do not have any direct mechanistic effects on community structure, but rather influence local factors that, in turn, will causally impact communities. Other regional factors, however, such as land-use cover are likely to have both direct (e.g., changes in habitat structure) and indirect (e.g., changes in water chemical quality) impacts on aquatic communities. Thus, local factors that may seem less important at first might effectively act as transmission belts, propagating a part or the total effects of some regional factors on community structure. Those effects are then likely to propagate within biological communities as a function of biotic interactions (e.g., effects on preys, which in turn, affect predators). Overall, we cannot rely on whole-community endpoint biodiversity measurements only, such as local species richness, to understand the direct and indirect pathways by which regional and local factors interact and propagate through biological communities to influence their structure and function^25–27^.

Here, we disentangled the main spatial, environmental and anthropogenic drivers shaping stream macroinvertebrate trophic structure across an entire river catchment. Starting from abundance data from a Swiss-wide biodiversity-monitoring program we collected functional traits on each taxon to reconstruct the trophic structure of each local community for 364 sites covering the entire Swiss Rhine river catchment. Integrating data related to land-use change, local water chemical and physical properties, regional factors related to altitude and position along the dendritic network, we used an integrative path analysis framework to identify specific pathways by which factors interact across spatial scales to affect stream invertebrate trophic structure.

## RESULTS

Testing the main spatial, environmental and anthropogenic drivers of aquatic macroinvertebrate trophic structure along the Swiss Rhine river catchment we found that variations in relative (Figure 1,2) and absolute (Figure 3) abundances of each trophic group across the whole river basin was largely driven by altitudinal variations (Table 1 and Figure 2). In turn, altitude influenced several other regional and local scale factors leading to a complex array of direct and indirect pathways across spatial scales, eventually leading to landscape wide variations in trophic community structure (Figure 1 & 3)

**Figure 1.**
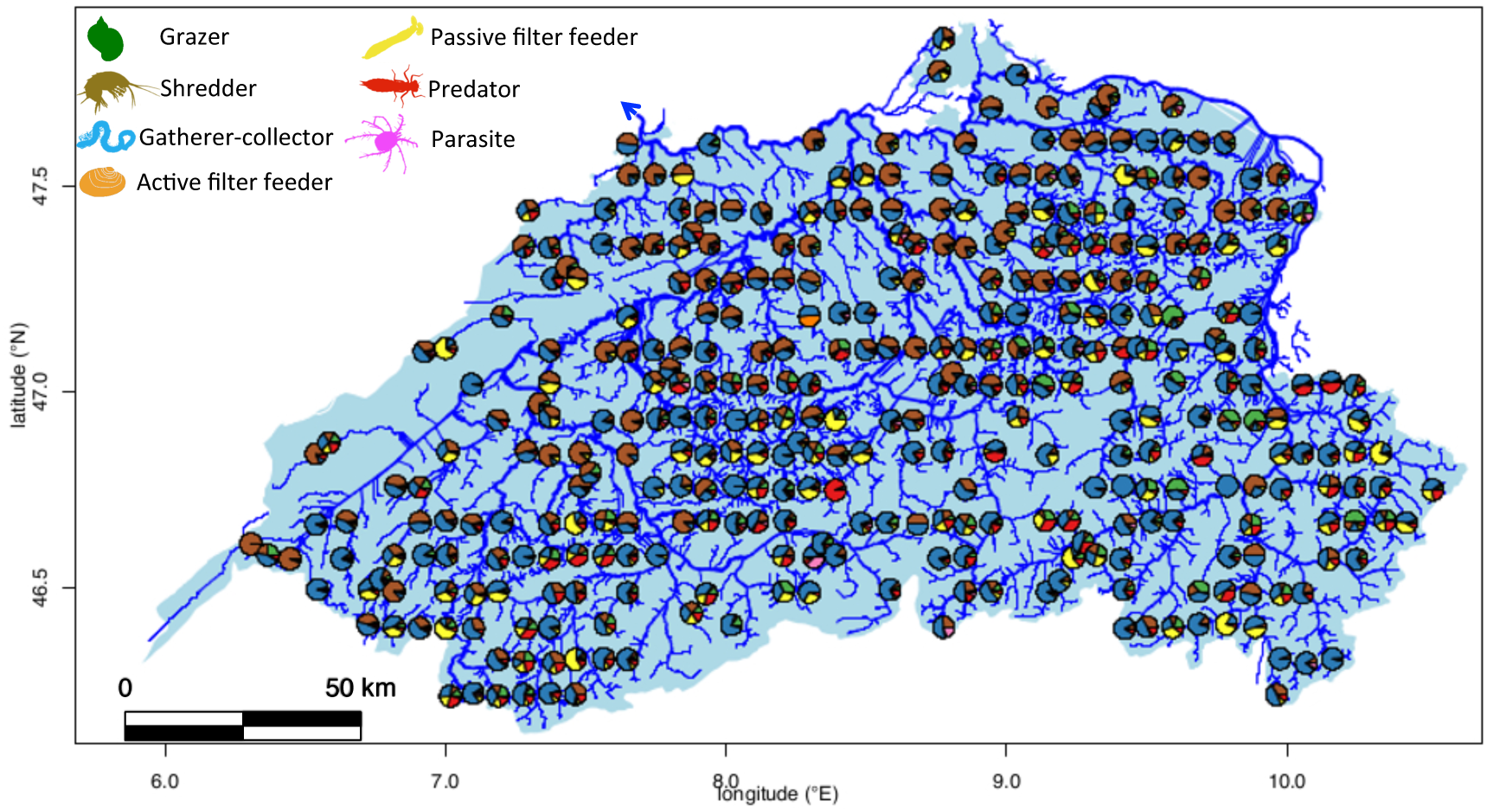
Spatial variation in trophic structure of riverine macroinvertebrates. The figure shows the Rhine river basin. All 3^nd^ order stream or larger are shown (arrow indicates direction of flows). Each pie chart represents the trophic structure (relative abundance of each trophic group in the community) for one of the 364 sampling sites across the river basin. Each functional group is represented by a silhouette of one of its iconic taxon: Gatherer-collector (Oligochaeta), Grazer-scraper (Limnaeidae), Predator (*Cordulegaster*), Passive filter feeder (Simuliidae), Active filter feeder (Sphaeridae), Parasite (Hydracaria), Shredder (Gammaridae).

**Figure 2.**
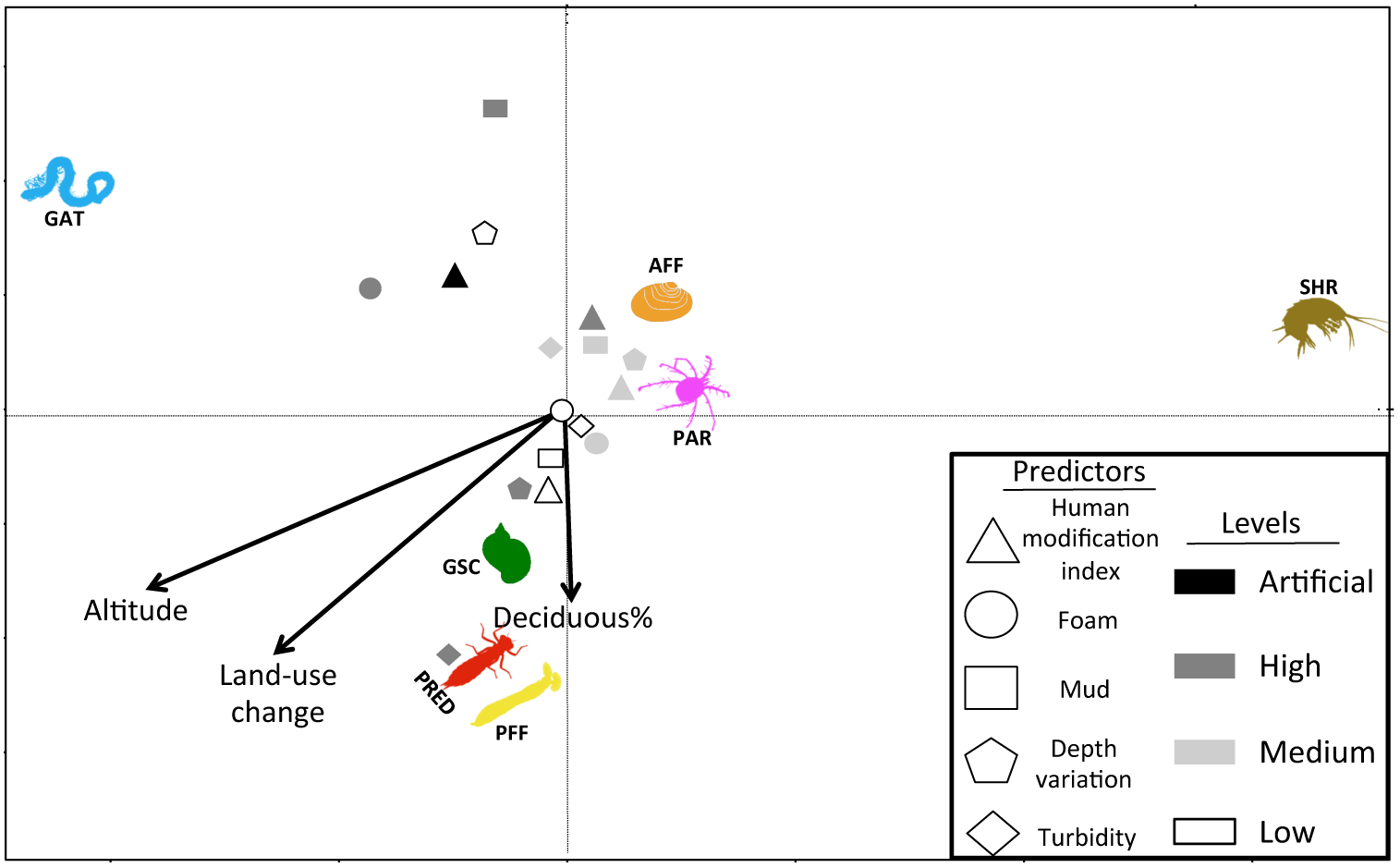
Main environmental and spatial drivers of riverine macroinvertebrate trophic structure. The ordination figure is the final db-RDA model selected by an automatic stepwise model building approach based on adjusted R^2^. The first and second axes respectively explain 67% and 18% of the total variation in trophic structure (relative abundance of each trophic group *per* community, see Methods). A specific geometric shape represents each categorical predictor, with the gray gradient representing the level of each predictor. Each functional group is represented by a silhouette of one of its iconic taxon: GAT: Gatherer-collector (Oligochaeta), GSC: Grazer-scraper (Limnaeidae), PRED: Predator (*Cordulegaster*), PFF: Passive filter feeder (Simuliidae), AFF: Active filter feeder (Sphaeridae), PAR: Parasite (Hydracaria), SHR: Shredder (Gammaridae).

**Figure 3.**
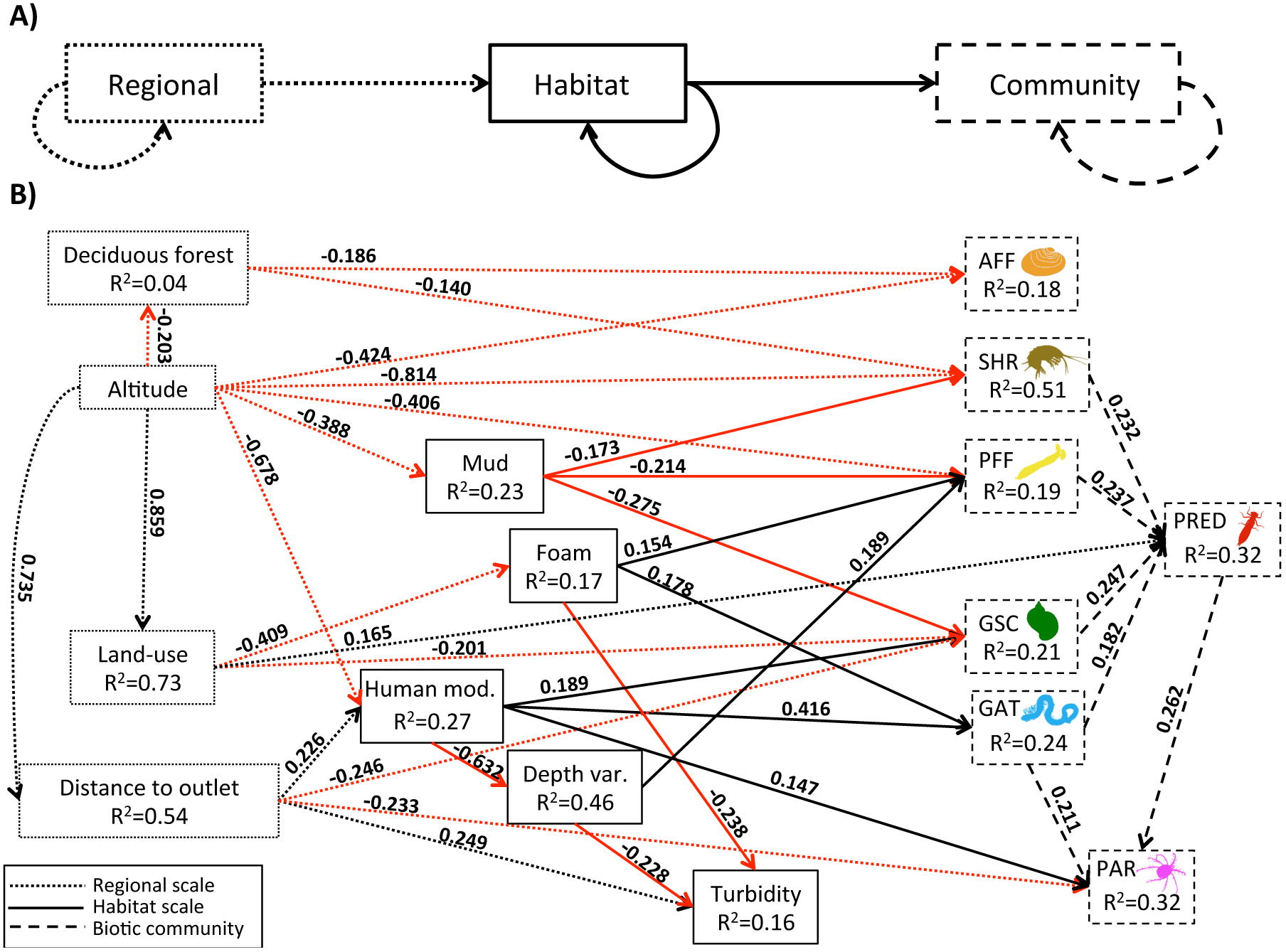
Direct and indirect pathways by which regional and local drivers influence riverine macroinvertebrate trophic structure. A) We hypothesized that most regional (dotted lines) factors would influence the biotic community (dashed lines) indirectly via an effect on local habitat factors (solid lines). We also expected within spatial scale interaction structure at both regional and local scales (looped arrows). Changes to biotic communities are usually analysed assuming that each predictor influence each taxa or functional group in the community, However we hypothesized that the specific structure of interactions within a biotic community would rather drive the propagation of effects from specific entry points (preys) to the entire community (looped arrow on the community box). B) Final structural equation model illustrating the different direct and indirect pathways by which regional (dotted lines) and local (solid line) factors interact and then propagate through the trophic community (dashed lines). Each value is the standardized coefficient (standardized estimate from each partial regression), representing the strength of the effect of one variable on another. Red arrows represent negative effects and black arrows positive ones. Each functional group is represented by a silhouette of one of its iconic taxon: GAT: Gatherer-collector (Oligochaeta), GSC: Grazer-scraper (Limnaeidae), PRED: Predator (*Cordulegaster*), PFF: Passive filter feeder (Simuliidae), AFF: Active filter feeder (Sphaeridae), PAR: Parasite (Hydracaria), SHR: Shredder (Gammaridae).

**Table 1.** Permutation ANOVA (200 permutations) on the final db-RDA model.

|  | Df | SS | F | *P*-value |
| --- | --- | --- | --- | --- |
| Altitude | 1 | 3.63 | 22.75 | 0.001 |
| Human Mod. Ind. | 3 | 2.16 | 4.51 | 0.001 |
| Foam level | 2 | 0.97 | 3.04 | 0.006 |
| Mud level | 2 | 0.88 | 2.75 | 0.004 |
| Land-use gradient | 1 | 0.81 | 5.10 | 0.001 |
| Depth variation | 2 | 0.95 | 2.98 | 0.012 |
| Deciduous cover | 1 | 0.60 | 3.80 | 0.011 |
| Turbidity level | 2 | 0.72 | 2.27 | 0.031 |
| Residual | 193 | 30.86 |  |  |

More specifically, altitude led to a decline in deciduous forest cover, was associated with an increase in distance to river outlet and drove land-use change from high settlement and agricultural lands to high altitudinal natural meadows (Figure 3). In turn, those regional factors influenced local habitats with transition to natural meadows leading to lower water foam levels (a proxy of eutrophication), and increased distance to outlet leading to higher turbidity level (Fig. 3). Lowland upstream sites were associated with higher probability of finding modified streams (see negative effects of altitude and positive effects of distance to outlet on river modification index on Figure 3). Local habitat factors then affected various trophic groups with mud level negatively impacting shredder, passive filter feeder and grazer-scraper abundances (Figure 3), foam (proxy of eutrophication) positively impacting gatherer-collector and passive filter feeder, river habitat modification positively influencing grazer-scraper, gatherer-collector and parasite, and higher riverbed variations in depth positively affecting passive filter feeder (Figure 3). Finally, all those regional and local factors affected predator abundance through affecting their preys (Figure 3).

We also found evidence for direct effects of regional factors on some trophic groups. Altitude had a direct negative impact on passive filter feeder and shredder abundances (Figure 3), potentially mediated by unmeasured (i.e., missing) local variables. Land-use transition to natural meadows had a direct positive effect on predator abundance suggesting that predators tend to fare better in high altitudinal streams surrounded by natural meadows than in low altitude zones characterized by a matrix of agricultural lands and human settlements, and a negative effect on grazer-scraper (Figure 3). Distance to outlet directly influenced gatherer-collector and parasite abundances negatively (Figure 3), illustrating that location along the river network has both indirect (mediated by local habitat factors, see above) and direct effects on aquatic invertebrate trophic structure.

There were also causal pathways among variables at each spatial scale as described above for altitude and regional factors. Among local factors, river modifications negatively impacted stream depth Variation, which in turn, negatively influenced turbidity (Figure 3). Thus river modifications had an indirect positive effects on turbidity mediated by a change in riverbed depth variation.

Overall, our results illustrate the complex interactions among local and regional scale predictors in shaping trophic structure along an entire catchment (Figure 3) and how the outcome of those interactions across-scale generate the specific context in which each trophic group will strive or not, leading to large spatial scale variations in trophic community structure (Figure 1 and 2).

## DISCUSSION

Testing for the main environmental and spatial drivers of trophic structure in stream macroinvertebrates we found a complex array of direct and indirect pathways by which regional and local drivers interact to influence relative and absolute abundances of aquatic macroinvertebrate trophic groups, eventually leading to landscape wide variations in trophic community structure. More specifically, cascading effects across spatial scales starting with altitude as a key driver influencing other regional factors, which in turn affected various local habitat characteristics directly to influence trophic group abundances. Most effects propagated through the community by first affecting preys, which in turn, affected predator abundances.

The importance of the river network has been shown to be context-dependant as a function of location-specific conditions such as terrestrial land-use and biotic interactions^14^. Our results suggest that those location-specific conditions can, in part, interact with some river network properties because they are not distributed randomly along the network but rather located at specific substructures in the network. Those effects constitute in themselves indirect effects of the river network rather than the absence of effect. More specifically, we showed that distance to outlet affected trophic groups directly but also indirectly via its positive effect on the human modification index. A main component of this result is the observation that lowland headwater locations are systematically more affected by human-induced riverbed and riverbank modifications than headwaters at higher elevations. Consequently, our results illustrate how the significance of spatial and regional factors can be masked by location-specific conditions when indirect pathways are not being taken into account^28^.

Our results also emphasize the complex response of each individual trophic group (see Figure 3) to each individual environmental and spatial factor. Interpreting any of these patterns independently can thus be misleading and only an integrative approach allows a coherent understanding of community structure, and eventually predicting shifts in response to multiple environmental changes^24,28,29^. Although our predictors are hierarchically organized (e.g., regional factors influencing local factors influencing preys, which in turn impact predators) rather than multiplicative, our study echoes recent calls to take a more integrative approach to the study of multiple-stressors and environmental changes, especially in aquatic ecosystems^25, 30–32^.

Shifts in trophic structure are a well-known driver of ecosystem processes^33–36^. Predicting those shifts, however, is a challenging endeavour because of multiple stressors interacting at different spatial scales and potentially affecting different trophic levels simultaneously. Our results suggest that effects mainly spread from preys to predators across the whole river network, and we observed important shifts in trophic group’s relative abundances. For instance, our ordination analysis identified an important gradient from gatherer-collector-dominated to shredder-dominated communities (see Figures 1 & 2). This observation is also visible with the structural equation modeling where higher gatherer-collector abundance is mainly associated with high levels of riverbed and bank modifications, while shredders seem to strive in less disturbed environments (Figures 2 & 3). At the functional level, we postulate that this shift from coarse (shredder) to fine particle (gatherer-collector) feeders along those environmental gradients is linked to variations in the type of resource available^37^. Such shifts in trophic structures have also implications for energy transfer and stoichiometric constraints in the community because shredders mainly feed on allochthonous leaf particles, which tend to be rich in carbon but nitrogen poor, while fine particles associated to agricultural lands tend to be nutrient rich but a poorer source of carbon.

Looking at the functional or trophic structure of communities is an essential step to better understand the effects of environmental perturbations on biodiversity and ecosystem functions, but also to eventually better predict the strength and direction of those effects^26,27,38^. Our results illustrate the complex interactions among local and regional scale predictors in driving trophic structure and how the outcome of those interactions across-scale generate the observed large scale variations in aquatic trophic community structure.

## METHODS

### Data

Our study used the aquatic macroinvertebrate abundance data from 364 sites across the whole Rhine river catchment in Switzerland, covering about 30,000 km^2^ and eventually flowing into the North Sea. The data is collected and curated by a Swiss governmental monitoring program (“Biodiversity Monitoring in Switzerland BDM”; BDM Coordination Office, 2014). Sampling is done following a systematic sampling grid, and was conducted in wadeable streams, 2^nd^ order or larger in size, thus excluding standing waterbodies, 1^st^ order streams and large rivers inaccessible by wading^39^. Each site was sampled once between 2009–2014 with seasonal timing of sampling adjusted with respect to elevation: the sampling period for a site was based on local phenology so as to collect as many macroinvertebrate taxa as possible for a given elevation^39^.

The survey was done using a standard kick-net (25 x 25 cm, 500μm mesh) sampling procedure defined in the Swiss “Macrozoobenthos Level I” module for stream benthic macroinvertebrates (BDM Coordination Office, see ^39,40^). Briefly, a total of eight kick-net samples were taken at each site to cover all major microhabitats within a predefined section of the river (area covered per site was width x 10 times the average width in length). Therefore, all locally represented habitat types (including various sediment types such as rocks, pebbles, sand, mud, submerged roots, macrophytes, leaf litter and artificial river-beds) and water velocities were sampled. Samples were preserved in 80% ethanol and returned to the laboratory for processing. In the laboratory, all macroinvertebrates used in this study were sorted and identified to the family level by trained taxonomists (total of 63 families see Table S1 for a list). For further details on the sampling method and the database, see also ^40–42^.

### Predictors

We used 38 predictors representative of regional, local and hydrological conditions, as well as land-use coverage and position in the dendritic network (see Table S2 for a complete list of each variable with description). Regional predictors included altitude at the sampling site and catchment size. Local predictors represent instream habitat conditions that were measured directly at sampling site. Local predictors included features of channel cross-section (e.g., width, depth, and their variability), riverbed conditions (e.g., mud deposition and attached algae), aquatic conditions (e.g., turbidity and dissolved iron sulfide concentration), and a discrete ranking of human alterations to riverbank and riverbed (see ^39^ for details). Hydrological predictors are factors representing geometry conditions of the river network in the upstream catchment of a sampling site. Those predictors included geomorphological (e.g., riverbed slope), hydrological (e.g., mean discharge) and chemical (e.g., inflowing wastewater volume) conditions. Land use predictors represent terrestrial conditions surrounding a sampling site. Those predictors included 6 land use classes considering adjacent influences to the local site with a lateral buffer distance of either 500 meters, 1, 5, 10, 100 or 1000 kilometers^42^. We know from previous work on this data that the 5 km scale is most significant in affecting stream invertebrate diversity^24^, thus we used only the 6 land use classes with lateral buffer distance of 5 kilometers in our analyses. Network predictors represent the position of each sampling site in the river dendritic network (e.g., centrality and distance to the outlet).

Many land use predictors were strongly skewed toward zero leading to important loss of information and degrees of freedom when analysing each variable individually. Instead, to emphasize a more continuous transition between each land-use type, for further analysis, we used scores from a canonical correspondence analysis representing a gradual shift in land-use from high proportion of human settlement and agricultural lands to high proportion of natural meadows (see Fig. S1). Such gradient is dominant in Switzerland with low lands representing most of the urban and agricultural lands. Grouping our land-use data this way reduced our total number of predictors to 34 for 364 sites.

### Trophic structure

We built the trophic structure of each stream macroinvertebrate community for each site, using the’freshwaterecology’ European database^43^ and extracting the’feeding type’ metric (*sensu* ^44^) for each of our 63 stream macroinvertebrate families. The data from the’freshwaterecology’ database was at the species level. Thus, we used averaged values across all species within family to determine the dominant feeding type of each 63 family. At the end, our data was comprised of abundance data for 63 families across seven functional feeding groups (following definition by ^44^, see Table S1) defining overall trophic structure. The seven groups were: grazer scrapers (13 families, mainly feeding on particulate organic matter from endolithic and epilithic algal tissues and biofilm), shredders (10 families, mainly feeding on coarse particulate organic matter from fallen leaves and plant tissue), gatherer collectors (10 families, mainly feeding on sedimented fine particulate organic matter), active filter feeders (1 family, mainly feeding on suspended particulate organic matter actively filtered from the water column), passive filter feeders (2 families, mainly feeding on suspended particulate organic matter passively trapped from running water), predators (24 families, mainly feeding on preys), and parasites (2 families, mainly feeding from hosts).

### Analyses

#### Ordination

To identify the main environmental and spatial drivers of the trophic structure of stream macroinvertebrate communities we used a distance-based redundancy analysis on Euclidean distances (db-RDA, following ^45^) followed by an automatic stepwise model building approach for constrained ordination based on the adjusted R^2^ of the full model (499 permutations, following ^46^). The significance level at P<0.05 of the final model, and of each selected term were tested using a permutation ANOVA (200 permutations). The uses of pairwise Euclidean distances ensure that our analyses really emphasize changes in the relative proportion of each trophic group within each community rather than between site changes in absolute abundance or composition^47^. Because we did not have any *a priori* knowledge on which predictors might be most important, we used all 34 predictors into our analytical pipeline. At the end 8 predictors were selected with first and second axes respectively explaining 67% and 18% of the total variance for the constrained axes.

#### Structural equation model

Ordination approaches provide insightful information on main drivers, however they do not provide information on the potential interactions and pathways by which each driver affect different trophic levels. For instance, a regional factor such as altitude does not have any direct ecological relevance. Rather, altitude will affect trophic groups via its effects on local factors (e.g., temperature or deciduous forest cover). Thus, even variables that may seem less important at first might act as transmission belts for the effects of other factors on stream invertebrate trophic structure. Moreover, factors affecting predators can do so by affecting the predator directly (e.g., high turbidity decreasing hunting efficiency) or indirectly by affecting its preys. Based on the information from the db-RDA analysis, we built a meta-model representing the potential links of importance in the system and how they affect trophic structure. We hypothesized that effects would mainly cascade from regional factors affecting local factors which in turn affect different trophic group (Fig. 1a). We then used structural equation modeling to test the fit of this initial meta-model against the data. Subsequently, we used the residual co-variance matrix and modification indices^48^ to identify potentially important missing links that were not included in the original meta-model. After adding those links to the model we then identified and pruned least important links (based on p-values and effect on model fit) to avoid over-parameterization and over estimation of explanatory power. Because we used categorical factors, we measured the fit of our model to the data with a robust diagonally weighted least square estimator (DWLS, see ^48^). Our final model converged after 105 iterations and showed a good fit to the data (n = 364, DWLS = 63.36, Degree of freedom = 62, P = 0.428).

All analyses were conducted with R 3.1.2 (R Core Team 2016), using the’vegan’ package^49^ for the db-RDA (‘capscale’ function) and stepwise model building (‘ordistep’ function), the’igraph’ package^50^ to compute network metrics, and the’lavaan’ package^48^ for SEM analysis.

## Acknowledgement

The Swiss Federal Office for the Environment (BAFU) provided the BDM data. We thank Roman Alther for discussion and for help in extracting GIS data. We thank the many people who conducted field and laboratory work within the biodiversity monitoring programme, and Nicolas Martinez and Tobias Roth for help in data provisioning. Funding was provided by the Swiss National Science Foundation Grants PP00P3_150698 and PP00P3_179089 (to F.A.).

